# Serum lipid reference intervals of high-density and low-density lipoprotein cholesterols and their association with atherosclerosis and other factors in Psittaciformes

**DOI:** 10.1101/2024.12.03.626694

**Authors:** Matthias Janeczek, Rüdiger Korbel, Friedrich Janeczek, Helen Alber, Helmut Küchenhoff, Monika Rinder

## Abstract

The prevalence of atherosclerosis is high in captive psittacine populations and the disease and subsequent heart problems have become a common reason for consultations in avian veterinary practices. To this day, ante-mortem diagnosis in birds remains challenging, however the serum lipoprotein-panel has been suggested to potentially aid in the diagnosis of the disease and provide better understanding of the pathogenesis. In parrots, unlike in humans, an association between serum lipoproteins and atherosclerosis has not been proven so far. The present retrospective cohort study aimed to establish multi-genera serum reference intervals for high-density-lipoprotein cholesterol (HDL-C) and low-density-lipoprotein cholesterol (LDL-C) in various parrot species. In addition, an evaluation on the changes of HDL-C and LDL-C depending on intrinsic and extrinsic factors of genus, age, sex, diet, prevalence of atherosclerosis, reproductive activity and body condition score was performed. An analysis of 1199 blood samples originating from 694 birds of multiple parrot genera revealed genus-specific variations in lipoprotein levels. Lipoproteins were studied for their association with independent variables of diet, age, sex, reproductive and breeding status, atherosclerosis and body conditioning score. A significant association between LDL-C levels and the prevalence of atherosclerosis could be observed, similar to findings in humans. Diet was found to be influencing both lipoprotein levels and risk for the development of atherosclerotic disease. Results showed, that LDL-C appears to be a potential predictor of atherosclerosis, while the role of HDL-C remained less conclusively defined. The results of the study provide a foundational framework for the future use of lipoprotein analysis in parrot medicine, offering novel insights into the management of cardiovascular health in pet parrots.

## Introduction

Parrots have been kept as companion animals for centuries with an increase in popularity in the late 20^th^ century (1). With time we have gained a better understanding of their biology, husbandry and disease patterns (2). Infectious diseases in parrots like psittacosis and psittacine beak and feather disease were once the focus of medical intervention for avian veterinarians worldwide. The parrot population in the western hemisphere that originated from large-scale imports in the late 20^th^ century (3) grew older and with rising age, avian veterinarians were confronted with importance in management of poor husbandry, and emergence of geriatric diseases.

Parrots are a zoological order of long living birds with the genetic potential to reach ages documented up to 88 years (4), depending on genus, species, as well as proper diet and husbandry. With progressing age of parrots, avian veterinarians were confronted with similar geriatric diseases as in humans, as older birds often developed arthrosis, cataracts and cardiovascular disease (5). Atherosclerosis in particular has become the cardiovascular disease with the highest prevalence in captive parrot populations with documented prevalences up to 91.8%, and is a commonly reported secondary finding in post mortem examinations (6–9). Atherosclerosis is generally characterized as a chronic inflammatory disease of the arterial wall (10). The accumulation of foam cells, lipids, calcium and cellular debris leads to atheromatous plaques that narrow the arterial lumen. In parrots, the disease itself often remains in a subclinical stadium and difficult to diagnose in its initial stage (7, 11). The ante-mortem diagnosis is commonly based on the emergence of actual clinical symptoms (12). However, even in that stage diagnosis is oftentimes difficult, as diagnostics often have low sensitivity and specificity, are high in cost or invasive (7, 11, 13–17). Such diagnostic measures include radiography, computed tomography, echocardiography, fluoroscopic angiography, blood pressure measurements, endoscopy or nuclear imaging (14), but to this day a standardised diagnostic protocol with high sensitivity and specificity remains to be established.

In humans in the western world, atherosclerosis and subsequent ischemic heart disease is the main cause of death, followed by strokes. Given its high clinical relevance, atherosclerosis has been the focus of extensive research, in which animal models played a key role (18), leading to a well-established understanding of its pathogenesis, clinical course, and therapeutic interventions. Influenceable predisposing factors were identified and found to include a sedentary lifestyle, obesity, diabetes mellitus, smoking, and many others, with hypercholesterolaemia, high low-density lipoprotein (LDL) and low high-density lipoprotein (HDL) being key factors in the development of atherosclerosis (19, 20). Lipid lowering drugs, primarily statins, are a main focus of therapy in humans with atorvastatin and rosuvastatin most commonly used in the treatment of dyslipidaemias and atherosclerosis (21). First studies about their pharmacokinetics, dynamics and potential therapeutic value in psittacines have been published (22–25).

Lipoproteins are molecular structures that consist of proteins and lipids, mainly cholesterol, triglycerides and phospholipids, that are formed in the liver of mammals, but also of birds (26). It is generally known that they play a crucial role in transporting lipids through the bloodstream to the peripheral tissues, where the lipids are used for energy utilization, lipid deposition, steroid hormone production and bile acid formation. HDL removes excess cholesterol from the bloodstream and tissues, transporting it back to the liver for excretion or recycling. LDL transports cholesterol from the liver to peripheral tissues. LDL consists of high amounts of cholesterol and few triglycerides, whereas HDL carries more protein and is of a higher density than LDL. Maintaining a balance between the various lipoproteins is essential for overall health. Dysregulation of lipoprotein metabolism may lead to various health problems, including atherosclerosis, heart disease, and metabolic disorders (10, 27). The opportunity to evaluate and properly understand additional lipoprotein data in HDL and LDL might further aid the diagnosis of dyslipidaemias and subsequent diseases like atherosclerosis, gonadal diseases, endocrine disease and nutritional diseases in parrots.

HDL and LDL are more correctly referred to as HDL-Cholesterol (HDL-C) and LDL-Cholesterol (LDL-C) in laboratory medicine, because not the concentration of HDL and LDL, but their cholesterol concentration in the serum is actually being measured. While testing blood chemistry values in clinical situations in parrot medicine, the lipid panel, typically consisting of total cholesterol (TC), triglycerides, HDL-C and LDL-C, is not commonly used. Though TC and triglycerides are sometimes evaluated, the literature lacks information on the diagnostic value of HDL-C and LDL-C, and there are few data on reference ranges for many different genera of parrots (28–30).

The goals of this study were as follows: First, to establish reference intervals for HDL-C and LDL-C serum concentrations in different parrot genera. Second, serum HDL-C and LDL-C concentrations were to be evaluated further, to understand, how the values shift with different environmental settings. More specifically, how the values change depending on age, sex, diet, body condition score (BCS), reproductive activity and actual prevalence of atherosclerosis. Third, birds with diagnosed atherosclerosis were observed based on the same factors, now including their individual HDL-C and LDL-C parameters, to evaluate the prevalence and risk factors associated with atherosclerosis in psittacine birds.

The hypotheses were that different parrot genera would have different reference intervals, and values would be higher for birds with unbalanced diets, reproductive activity, elevated body conditioning scores, atherosclerotic disease, as well as higher with rising age and in female sex. In addition, we expected birds would be more likely to develop atherosclerosis the higher their levels of LDL-C were and the lower HDL-C was. The same was hypothesized to be valid for rising age, female sex, elevated body conditioning score, unbalanced diet and breeding birds.

## Materials and methods

### Study animals

A total of n = 1196 HDL-C and 1190 LDL-C serum values were collected from n = 1199 blood samples from birds belonging to the zoological order Psittaciformes (generally referred to as parrots). The samples, collected over a period of seven years from November 2016 to September 2023, all originated from patients of a veterinary practice specialized and limited to parrots, living in Germany and the surrounding European region. A total of 14 different genera and 46 individual species were sampled.

Blood samples originated from leftover blood from different diagnostic workups of the individual bird in all cases. The birds were sampled due to various veterinary reasons, but not with the sole goal of obtaining HDL-C and LDL-C values. This project has been evaluated and approved by the ethics committee of the Veterinary Faculty of Ludwig-Maximilians-Universität München under AZ 376-25-09-2023. Reporting of this retrospective study followed the ARRIVE 2.0 guidelines (31). Birds were fasted for at least 6 hours prior to blood collection. Right jugular venipuncture was performed by standard measure (31) using a 26-gauge, 12mm needle attached to a 2 ml Syringe. The blood was collected into Serum Gel tubes (Sarstedt AG & Co. KG, Nümbrecht, Germany) and centrifuged 30 minutes after sampling. The serum was collected, cooled at 4 °C and tested at Antech Lab Augsburg, Germany (previously Synlab Holding, Augsburg, Germany) on an Alinity ci-series c-module (Abbott, Abbott Park, Illinois, U.S.A.) photometrically with an enzymatic colour test.

Additional information considered from anamnesis and physical examination of the birds included genus, species, age (if known), diet, body condition score, breeding condition and current reproductive status. Some patients presented at the practice for routine check-ups and consequently had their blood sampled, HDL-C and LDL-C measured and were included in this study more than once. Hence the 1199 blood samples originate from a total of 694 subjects consisting of 249 parrots from the *Amazona* genus, 88 from the genera *Ara* and *Anodorhynchus*, 18 from the genus *Pionites*, 8 *Pionus*, 231 *Psi*ttacus, 19 *Poicephalus*, 23 *Eclectus*, 3 *Ar*atinga, 2 *Deroptyus*, and 53 subjects from the family Cacatuidae (32). The sample pool for the genera of *Aratinga*, *Deroptyus* and *Pionus* were deemed too small for the quantitative analysis, therefore these birds were fully excluded from further evaluation. Sex was unrelated to this study determined by DNA-sexing (PCR) from blood or feather, surgical sexing via endoscopy, post-mortem examination or dimorphism in select species. Data were based on observations from 371 males and 323 females. The age of individuals ranged from 1 to 56 years. The birds’ diet was recorded and categorized in three categories. Category 1 included birds fed exclusively on seed mixtures, category 2 consisted of birds fed a mixture of seeds, food from table and some pellets or extrudates. Category 3 consisted of birds exclusively fed a pelleted or extruded diet, with the majority of the birds in this category fed Harrison’s bird foods High Potency or Adult Lifetime formulas (Harrison’s bird foods, Brentwood, Tennessee, U.S.A.). The intake of fruits and vegetables was not included in this variable. During physical examination a BCS of the individual bird was recorded from 1 – 5, with 1 being anorexic, 3 being normal and 5 being obese. The variable of breeding status was classified as a summary of the bird’s reproductive history, with three different categories. Category 1 included sexually immature, juvenile birds with no reproductive history. Category 2 included adult, sexually mature birds that have gone through breeding seasons and experienced the hormonal changes that come with them. Category 3 consisted of female parrots that have laid eggs in the past, regardless of them being fertilized or not. On the other hand, the variable of reproductive status defined the bird’s current reproductive activity at the time of sample collection, which was also grouped in different categories. This evaluation was made based on seasonal behavioural changes, copulation, changes to the birds’ cloaca under the influence of estrogen and endoscopic evaluation of the bird’s gonads (7). Category 1 included juvenile or reproductively inactive birds, due to factors like age or disease. Category 2 included adult birds outside of breeding season, showing no signs of reproductive behaviour, whereas birds that were under hormonal influence during the breeding season and exhibited said behaviour were put into category 3. Lastly category 4 was a category exclusively for adult, female parrots that laid eggs within one week before or after the time of sample collection.

Whenever possible, an additional screening for atherosclerosis was included, either by x-ray if severe calcification of the great vessels was visible, endoscopic evaluation of the large heart vessels and aorta descendens or post mortem examination in deceased birds (7, 17, 33). Due to a lack of an existing classification system for endoscopic evaluation, the observations were grouped into different categories, either with no data available, no atherosclerosis observed, mild atherosclerosis, moderate atherosclerosis or heavy atherosclerosis. The lesions were classified based on the size, overall number and colour intensity of the whitish-yellow discoloration of the arterious plaques. Because only 8 observations of heavy atherosclerosis were recorded, these birds were included into the pool for moderate atherosclerosis, to enable estimation of the statistical model.

All birds were evaluated as healthy or sick based on physical examination and individually available laboratory test results. Due to all birds being pets or kept in breeding facilities, no generalized categorisation for ambient temperature, air humidity or photoperiod was performed. The same is applicable for exercise and ability of flight, due to the huge differences and the lack of an existing categorization model for exercise.

For establishing the reference intervals, only clinically healthy birds without atherosclerosis or other signs for disease complexes and symptoms were included, to ensure a healthy reference population. Exclusion criteria varied greatly due to the variance of health problems in parrots, but included among others anaemia, leucocytosis, obesity, cachexia, pneumonia, cardio-vascular disease, hepatopathy, nephropathy, reproductive disease, egg laying and neoplasia. This left the reference intervals with 761 observations from 420 parrots in total consisting of 216 males and 204 females. Due to differences in the missing values between HDL-C and LDL-C, the according reference populations differed slightly. For the second aim of the study, all birds with existing HDL-C and LDL-C values, regardless of health status, were included. However, 313 observations with a missing diagnosis concerning atherosclerosis were excluded. Further, 14 observations with missing values in either one of the independent or dependent variables included in the three developed models for HDL-C, LDL-C and atherosclerosis, were excluded as well. This left 853 observations from 503 birds as the sample base for estimating the regression models. For analysing potential risk factors for atherosclerosis, a subset of the data without repeated measurements was used, to ease convergence of the model estimation. One subject was excluded from the analysis as it developed atherosclerosis during the repeated measurements. Furthermore, since none of the 12 *Pionites* birds exhibited atherosclerosis, all birds belonging to this genus were excluded from the analysis. This left the analysis with 490 birds and one observation each. A detailed overview showing how many birds were included in each model and the process of exclusion is stated in S1 Fig.

### Statistical analysis

All statistical analyses were conducted using R (v4.4.2; R Core Team 2024). For obtaining reference intervals for HDL-C and LDL-C, the empirical 5^th^, 50^th^ (median) and 95^th^ quantiles, as well as the total range of values, were calculated for the previously described reference subset of the base population.

To further investigate influences on HDL-C and LDL-C values, two linear mixed-effects models (LMM) were constructed, one for each blood value, and applied using the linear mixed-effects regression (lmer) function of the lme4 package (v1.1.35.5). Both models incorporated several independent variables, namely genus, age, sex, level of atherosclerosis, diet, current reproductive status and the body condition score. Additionally, random intercepts were included to account for individual variations among the birds. To assess the overall significance of the independent variables, Type II tests were calculated on the estimated models using the car package in R (v3.1.3) (34).

For investigating potential risk factors for atherosclerosis, the outcome variable was transformed into a binary variable indicating the presence or absence of atherosclerosis in birds. This transformation implies the application of a logistic Generalized Linear Model (GLM) to the data, here without using repeated measures. This was done using the glm function of the stats package (v4.4.2). The independent variables genus, age, sex, diet, breeding conditions and HDL-C and LDL-C levels were included in the GLM. As for the two LMM for HDL-C and LDL-C, a Type II test was conducted to estimate the overall significance of the independent variables. An alpha level of 0.05 was chosen as the threshold for statistical significance. The code and data used to conduct the analyses described can be found S1 and S2 files.

## Results

### Reference ranges of HDL-C and LDL-C

Blood concentrations for HDL-C and LDL-C obtained from the parrots included in this study are summarized in table 1 and 2 and figures 1 and 2. Genus related differences were clearly visible, meaning that different genera of parrots have different reference intervals of HDL-C and LDL-C and need to be evaluated accordingly in practice. The lowest median HDL-C values were recorded in macaws (genera *Ara* and *Anodorhynchus*), the highest in *Eclectus* parrots, followed by grey parrots (*Psittacus*) and amazons. For LDL-C, the lowest median was again observed in macaws, whereas was the highest was found in *Eclectus* and *Psittacus*.

**Figure 1.**
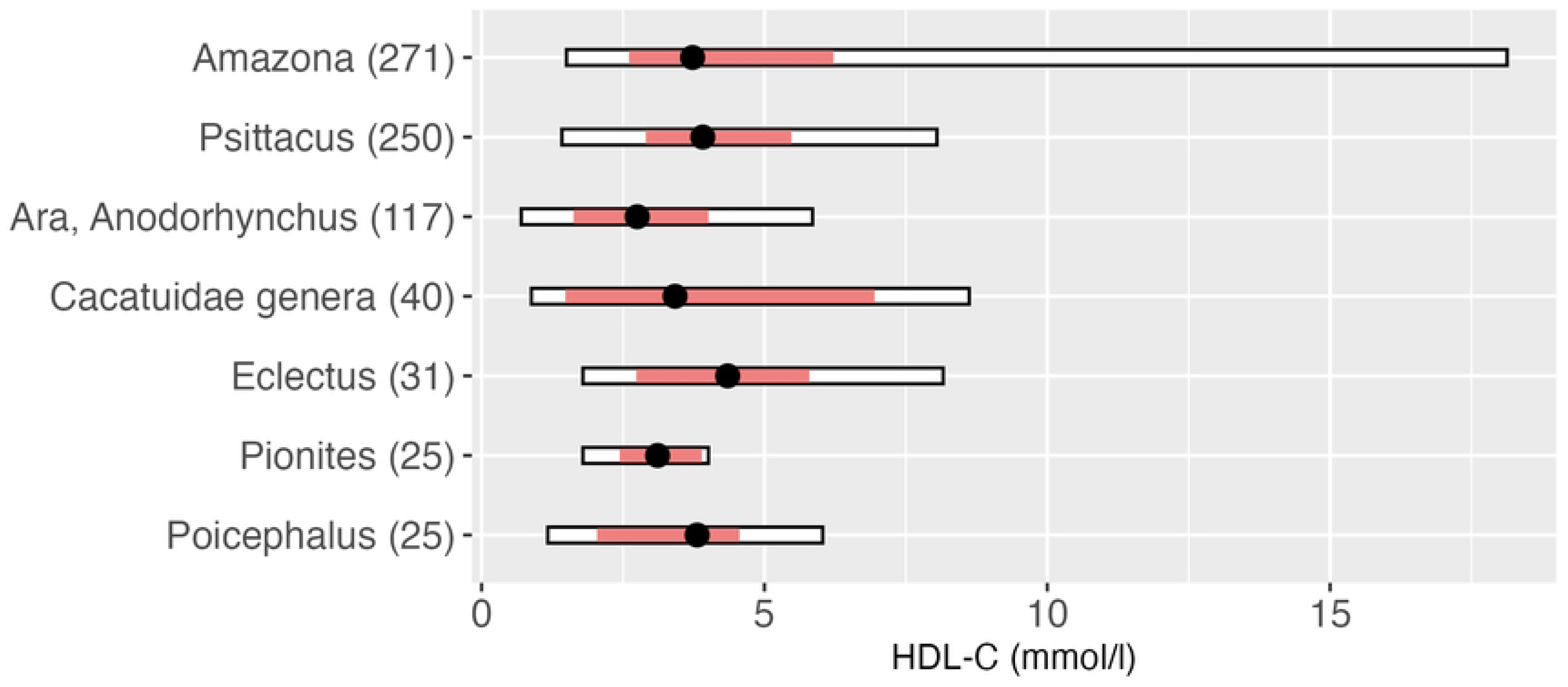
HDL-C reference values in mmol/l within genera giving the range, the median, the 5^th^ and 95^th^ percentiles, number in brackets behind genus indicates number of observations of the according genus

**Figure 2.**
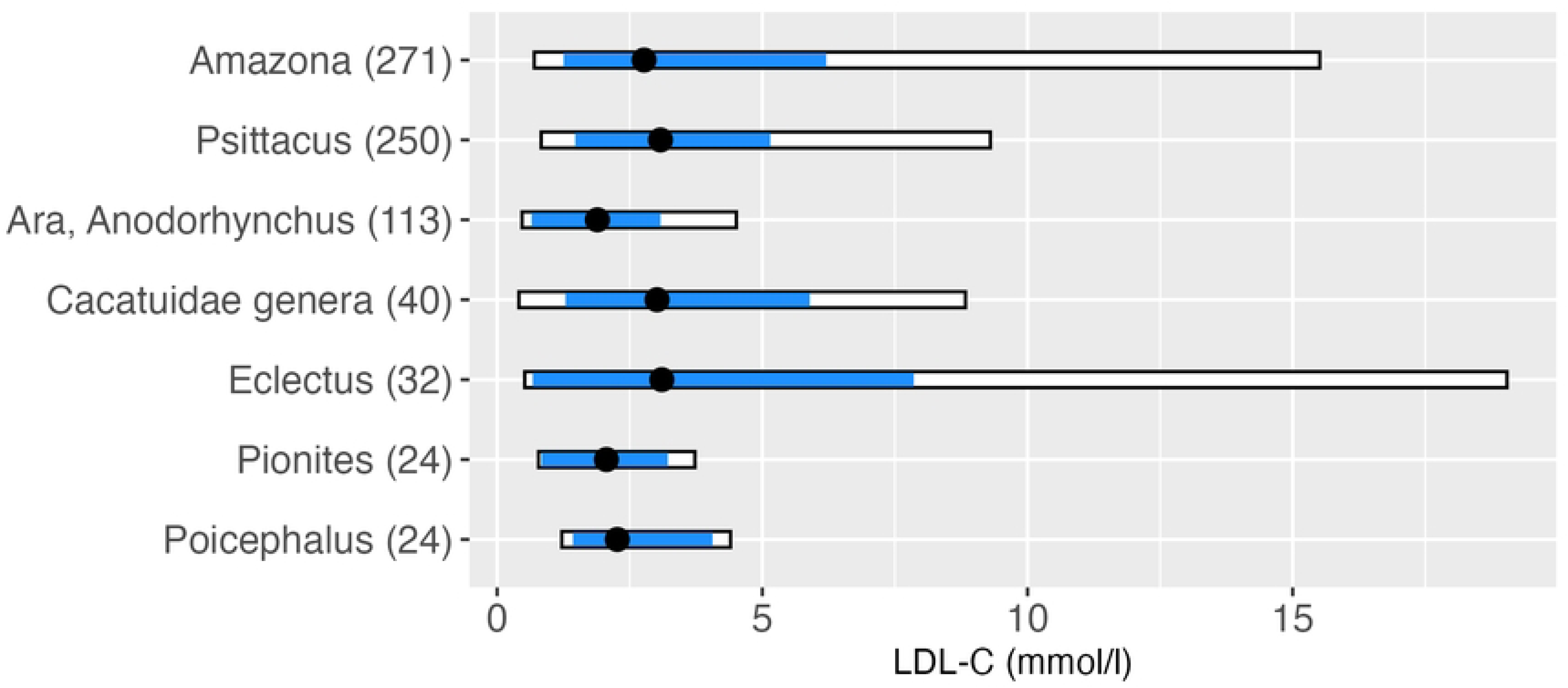
LDL-C reference values in mmol/l within genera giving the range, the median, the 5^th^ and 95^th^ percentiles, number in brackets behind genus indicates number of observations of the according genus

**Table 1:** Reference intervals for HDL-C blood concentrations (mmol/l) in select psittacine species.

| Genus | Sample Size | Mean | Range | Median | 5%-quantile | 95%-quantile |
| --- | --- | --- | --- | --- | --- | --- |
| *Amazona* | 271 | 4.04 | 16.6 | 3.73 | 2.60 | 6.22 |
| *Psittacus* | 250 | 3.99 | 6.63 | 3.91 | 2.90 | 5.48 |
| *Ara* and *Anodorhynchus* | 117 | 2.79 | 5.15 | 2.75 | 1.63 | 4.01 |
| Cacatuidae | 40 | 3.92 | 7.74 | 3.42 | 1.48 | 6.95 |
| *Eclectus* | 31 | 4.42 | 6.37 | 4.35 | 2.74 | 5.80 |
| *Pionites* | 25 | 3.20 | 2.22 | 3.11 | 2.44 | 3.89 |
| *Poicephalus* | 25 | 3.63 | 4.86 | 3.81 | 2.04 | 4.56 |

**Table 2:**
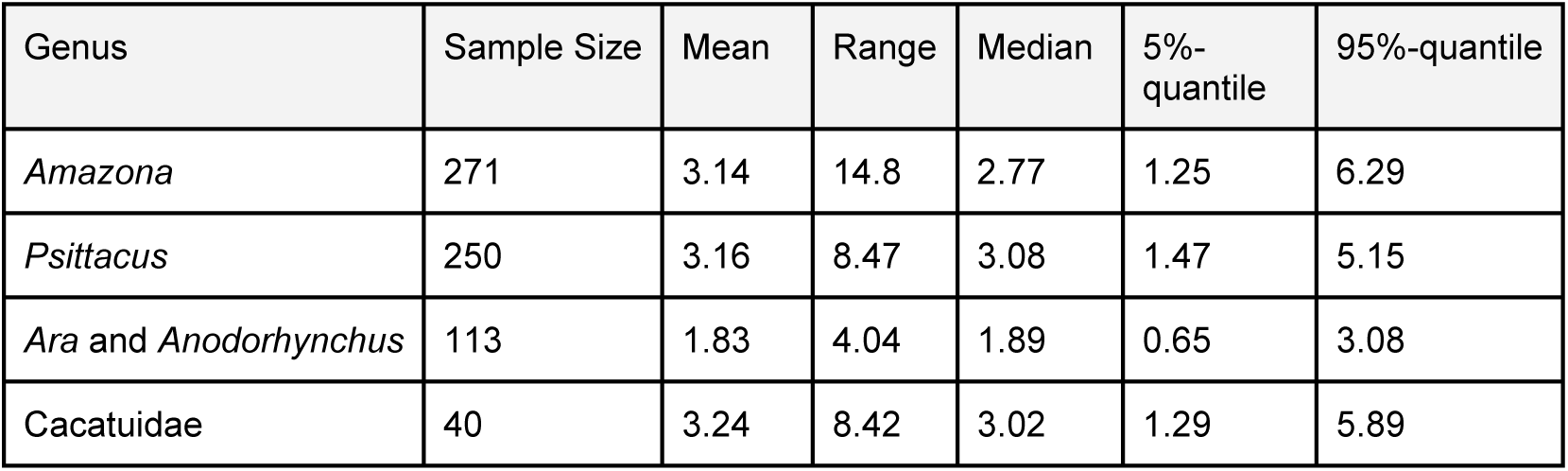
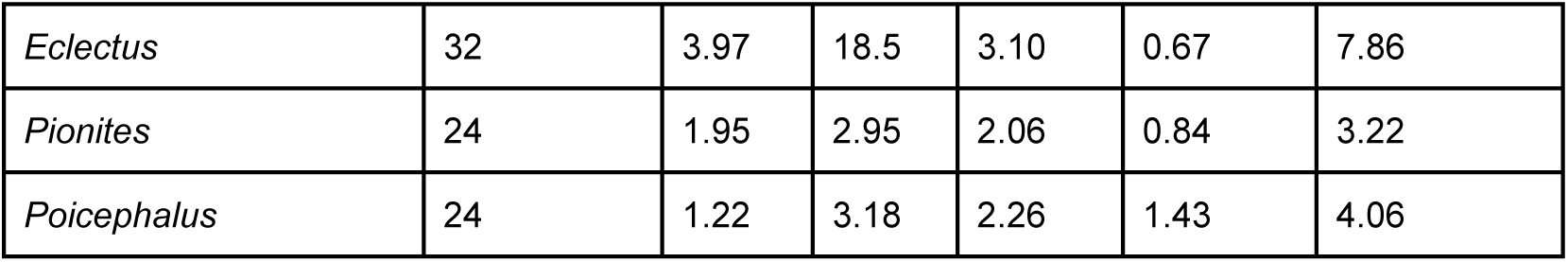
Reference intervals for LDL-C blood concentrations (mmol/l) in select psittacine species.

| Genus | Sample Size | Mean | Range | Median | 5%-quantile | 95%-quantile |
| --- | --- | --- | --- | --- | --- | --- |
| *Amazona* | 271 | 3.14 | 14.8 | 2.77 | 1.25 | 6.29 |
| *Psittacus* | 250 | 3.16 | 8.47 | 3.08 | 1.47 | 5.15 |
| *Ara* and *Anodorhynchus* | 113 | 1.83 | 4.04 | 1.89 | 0.65 | 3.08 |
| Cacatuidae | 40 | 3.24 | 8.42 | 3.02 | 1.29 | 5.89 |
| *Eclectus* | 32 | 3.97 | 18.5 | 3.10 | 0.67 | 7.86 |
| *Pionites* | 24 | 1.95 | 2.95 | 2.06 | 0.84 | 3.22 |
| *Poicephalus* | 24 | 1.22 | 3.18 | 2.26 | 1.43 | 4.06 |

### HDL-C model

Within the following, all effect estimates reported were to be interpreted on average and given all other variables in the according model were kept constant. Overall, the LMM applied reveals a significant relationship between HDL-C levels and several independent variables. Genus was found to be significantly associated with HDL-C levels (p < 0.001) (Fig 3), confirming the large differences between the genera of HDL-C values observed in the analyses of the empirical quantiles. The BCS also showed a significant effect on HDL-C levels (p < 0.001), with higher levels of BCS, BCS 4 (β = 0.41, SE = 0.16, p = 0.01) and 5 (β = 0.69, SE = 0.26, p = 0.01) being associated with increased HDL-C levels, and vice versa for the lowest BCS (β = -0.91, SE = 0.34, p = 0.01) being associated with lower HDL-C levels, as compared to HDL-C levels corresponding to a healthy BCS (Fig 4). No significant associations between HDL-C levels and age, sex, diet, reproductive status and atherosclerosis prevalence were observed (S2 Fig and S1 Table.).

**Figure 3.**
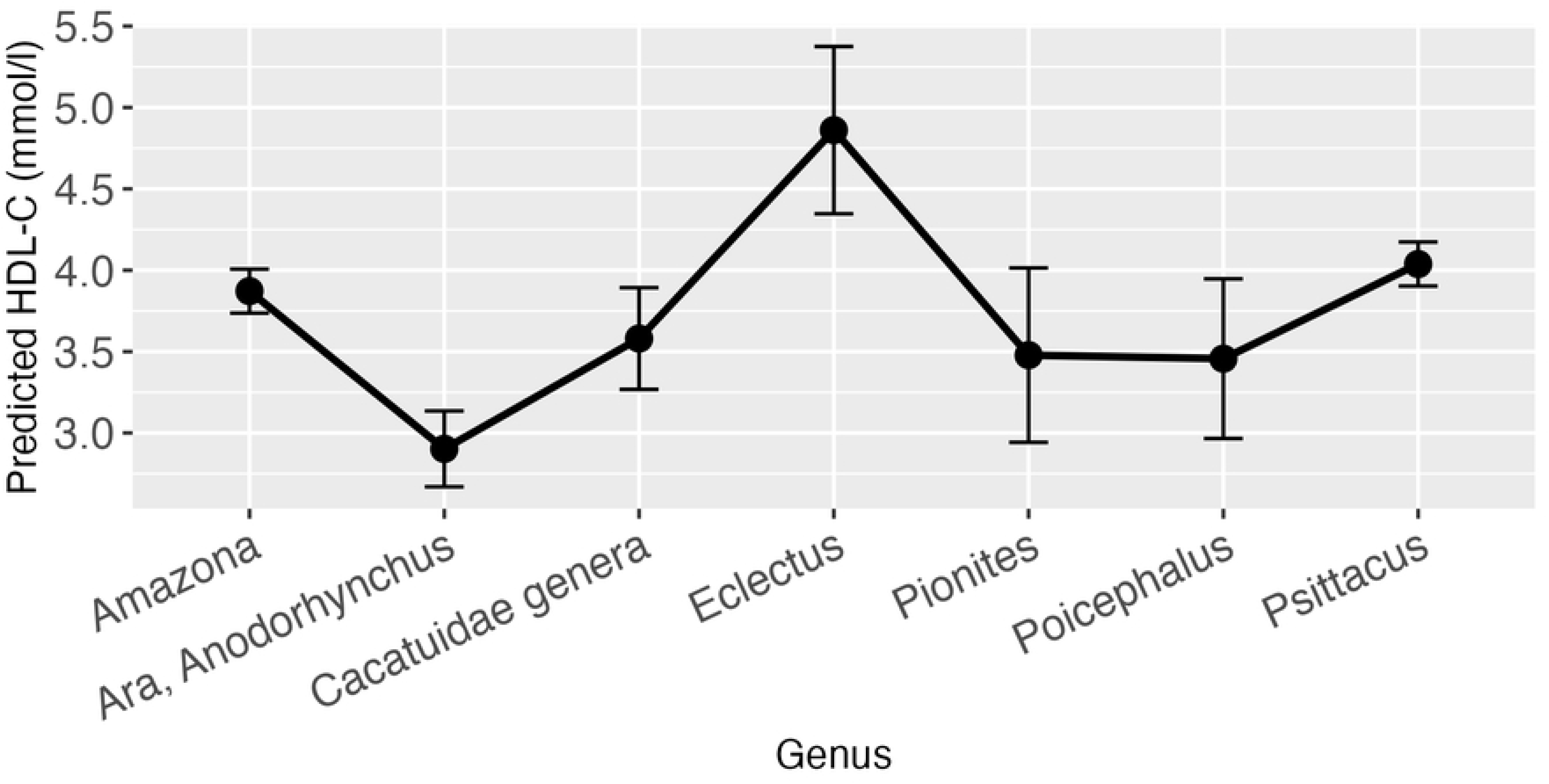
Effect plot with confidence intervals for HDL-C and genus.

**Figure 4.**
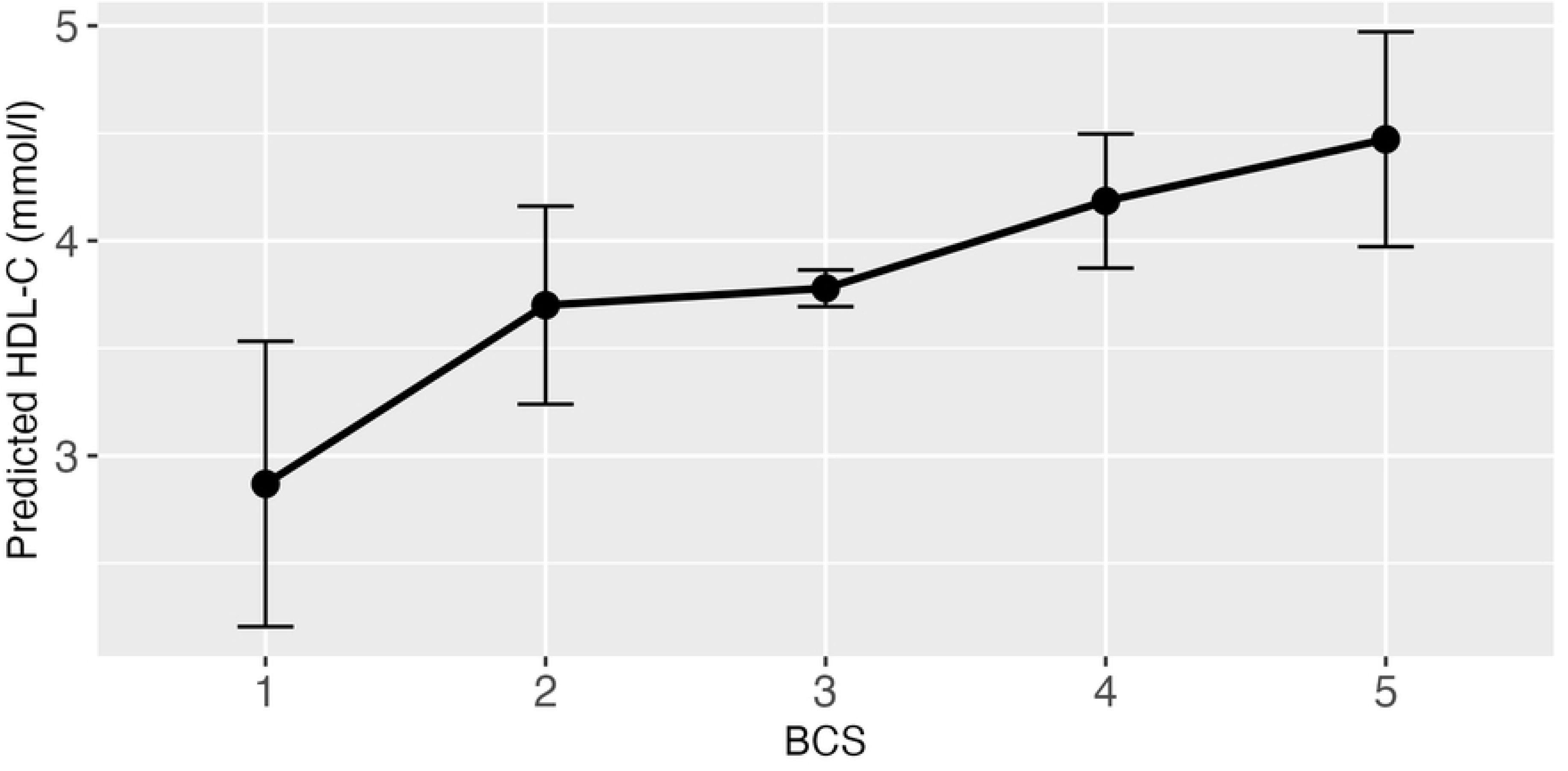
Effect plot with confidence intervals for HDL-C and BCS.

### LDL-C model

When investigating influences on the LDL-C levels, several significant relations could be identified through application of the LMM. As it was the case for HDL-C, there was a strong effect of the genera (p < 0.001) (Fig 5), confirming once again the large differences between birds from different genera as observed for the reference values. There was a significant relation between BCS (p = 0.03) and LDL-C values. The only significant effect on LDL-C within BCS, when compared to a healthy score, was observed for anorexic birds (β = 1.42, SE = 0.55, p = 0.01). However, it has to be stated, that for this data subset, only 9 observations were available for anorexic birds from which one value represented an outlier (LDL-C = 15.95 mmol/l). Neither the overall effect of the BCS nor the effect of the anorexic sample on LDL-C remained when this outlier was removed (Figures S3 and S4). Additionally for LDL levels, atherosclerosis (p = 0.001) as well as the diet (p = 0.03) displayed significant overall effects (Fig 6 and 7). More precisely, birds that were diagnosed with moderate to heavy atherosclerosis (β = 0.73, SE = 0.30, p = 0.01) seemed to have increased levels of LDL-C as compared to birds without atherosclerosis. This effect was not observable in birds diagnosed with mild atherosclerosis (β = -0.17, SE = 0.20, p = 0.38). Further, birds exclusively fed a pelleted or extruded diet (category 3) (β = -0.45, SE = 0.17, p = 0.01) displayed significantly lower LDL-C values as compared to birds exclusively fed on seeds (category 1), while there is no significant difference between birds from category 1 and birds fed a mixture of seeds, food from table and pellets or extrudates (category 2) (β = -0.31, SE = 0.21, p = 0.15). There appears to be a significant relation between reproductive status and LDL-C (p = 0.01), with the group of adult female parrots that laid eggs within one week before or after the time of sample collection (category 4), consisting of 10 animals, driving this main effect (β = -1.70, SE = 0.61, p = 0.01).

**Figure 5.**
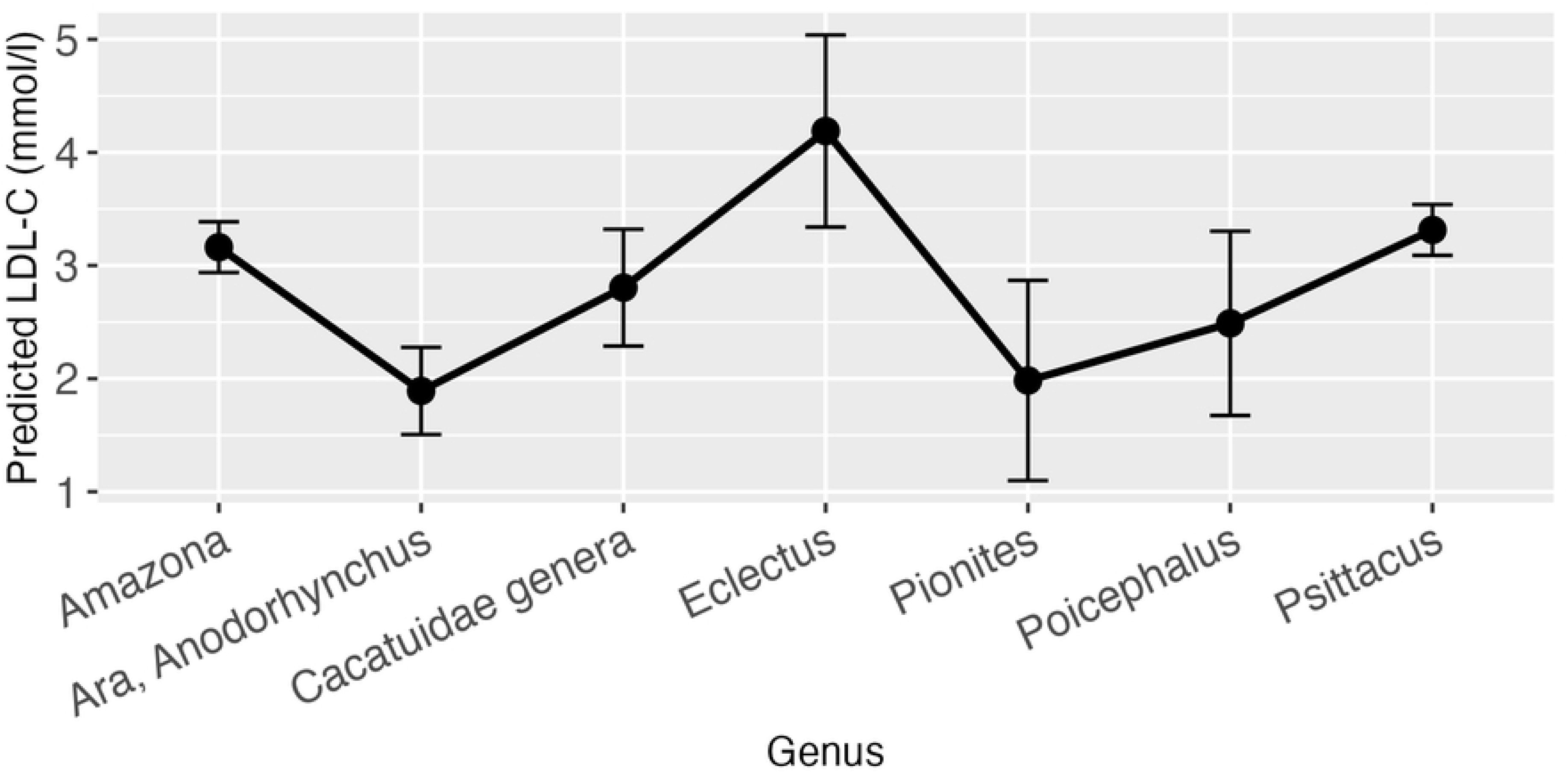
Effect plot with confidence intervals for LDL-C and genus.

**Figure 6.**
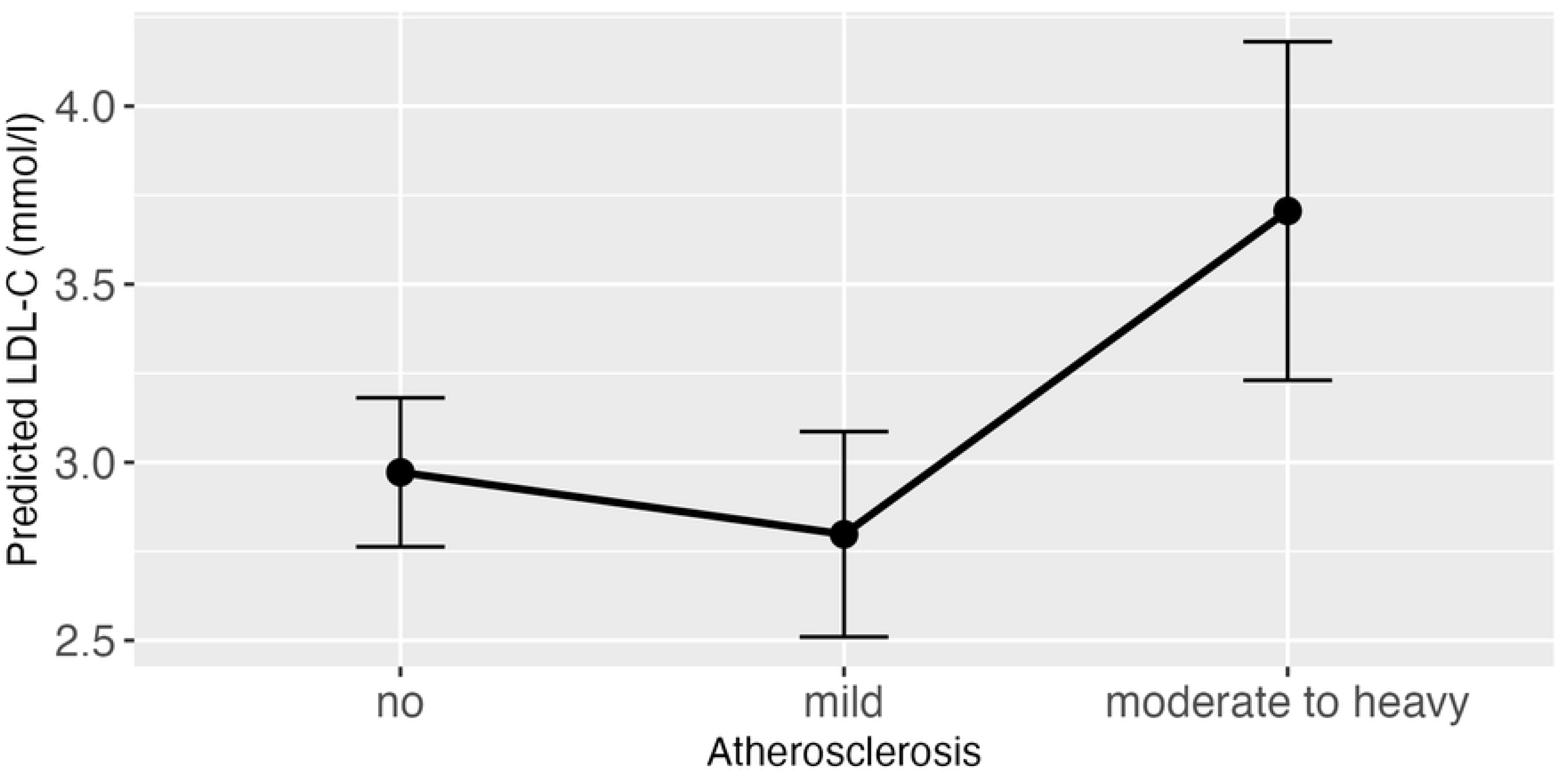
Effect plot with confidence intervals for LDL-C and atherosclerosis.

**Figure 7.**
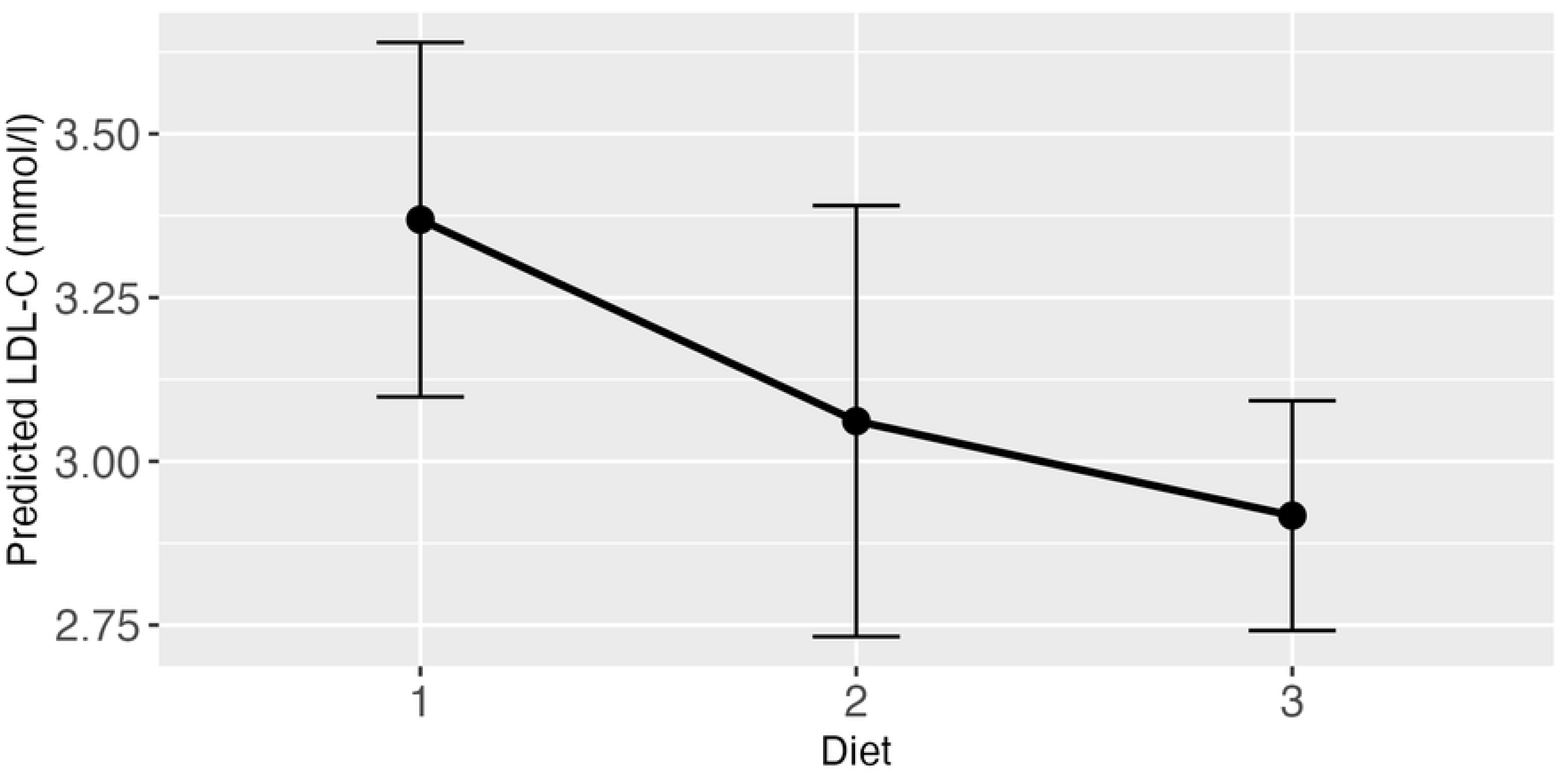
Effect plot with confidence intervals for LDL-C and diet. (category 1 pure seed diet, category 2 mixed diet seeds, table food, pellets/extrudates, category 3 pure pellets/extrudates)

### Atherosclerosis model

The GLM, modelling influences of various variables on the odds to be diagnosed with atherosclerosis, displayed several significant relations. Firstly, for a one-year rise in age, the odds of being diagnosed with atherosclerosis rose by a factor of 1.18 (β = 0.16, SE = 0.02, p < 0.001). Further, diet had a significant overall influence (p < 0.001), with birds fed according to category 2 (β = -1.18, SE = 0.40, p = 0.003) as well as birds fed according to category 3 (β = -1.73, SE = 0.32, p < 0.001) displaying lower probability for atherosclerosis as compared to birds fed according to category 1 (Fig 8). Additionally, a significant difference between the genera was found (p = 0.02) (Fig 9). No significant correlation between atherosclerosis and the variables sex, HDL-C, LDL-C, BCS and breeding condition were observed.

**Figure 8.**
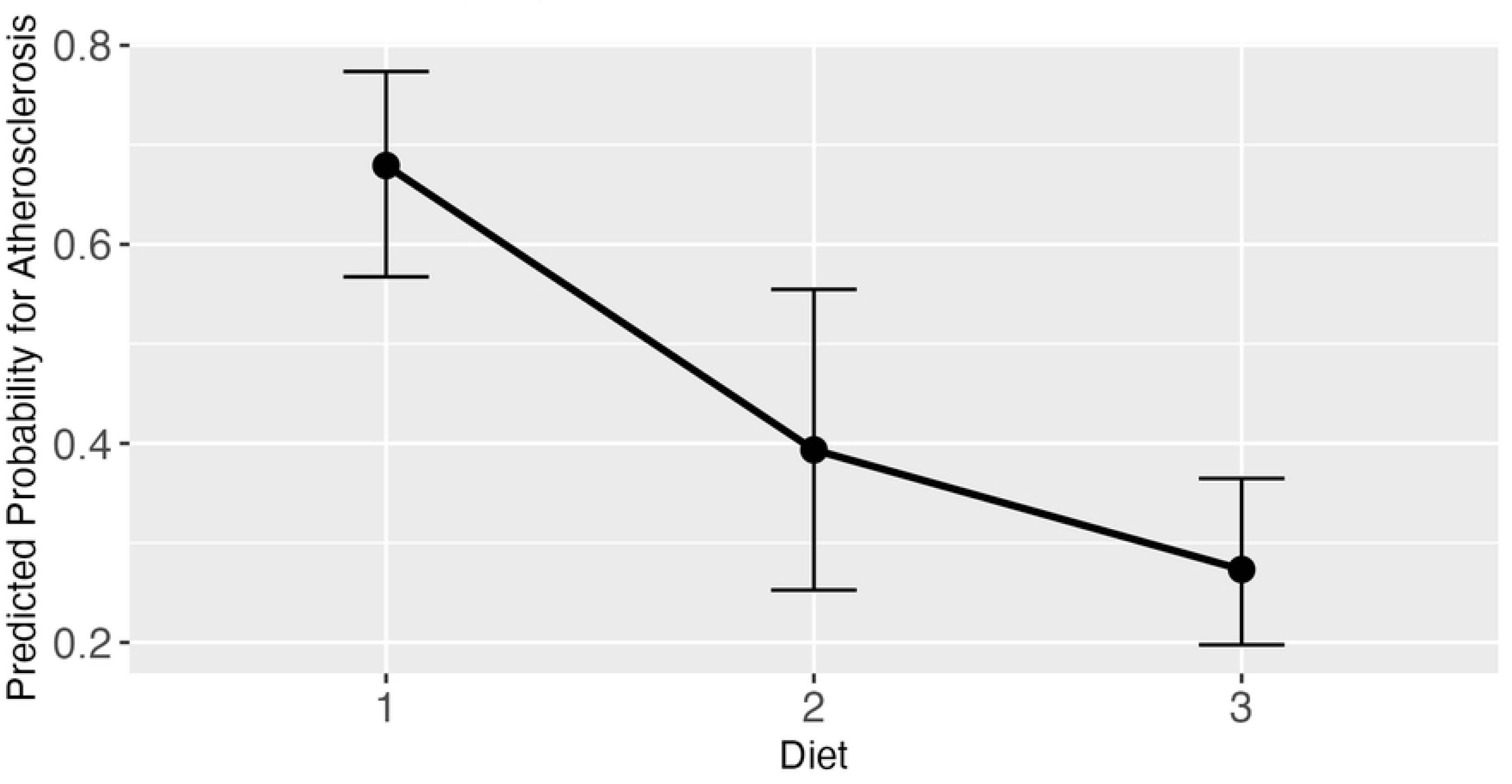
Effect plot with confidence intervals for atherosclerosis and diet. (category 1 pure seed diet, category 2 mixed diet: seeds, table food, pellets/extrudates, category 3 pure pellets/extrudates)

**Figure 9.**
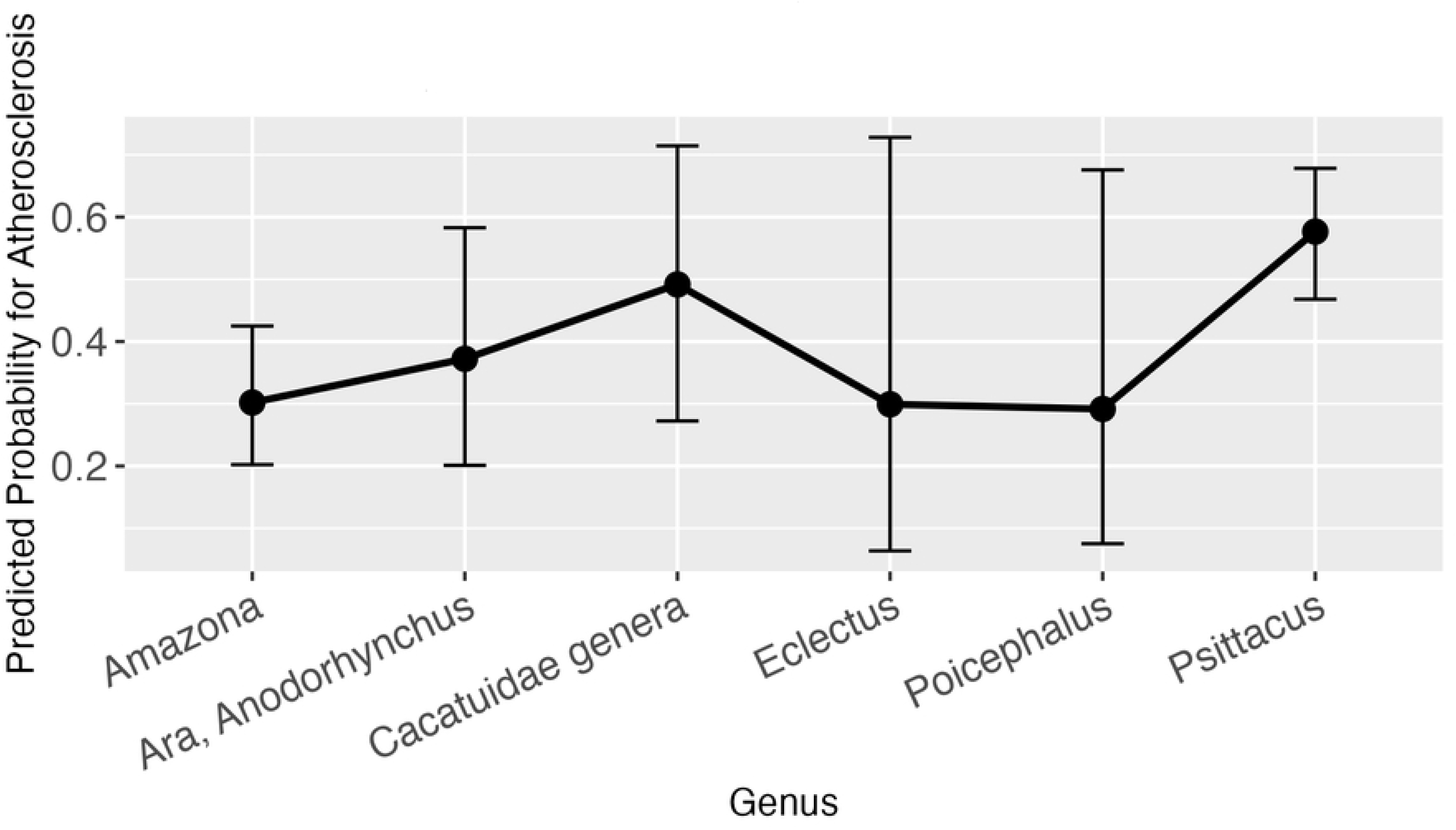
Effect plot with confidence intervals for atherosclerosis and genus.

## Discussion

The results of this study present large-scale, multi-genera reference intervals HDL-C and LDL-C in pet parrots. The analysis of reference intervals revealed significant differences associated with the genus. HDL-C and LDL-C values differed between genera and need to be interpreted accordingly. The genus *Ara*, including *Anodorhynchus*, which in the past have been reported to be the least susceptible genus to atherosclerosis (9), also showed the lowest levels of HDL-C and LDL-C. The genera of *Psittacus* and *Amazona*, which have been previously shown to be highly susceptible to atherosclerosis (9), showed very high median values for HDL and LDL. The genus of *Eclectus*, also commonly observed with atherosclerosis, showed high medians of HDL-C and LDL-C, however the incidence of atherosclerosis in *Eclectus* parrots has previously been suggested to be lower than in African grey parrots or amazons (12, 35). Additionally, *Eclectus* as a genus is difficult to evaluate, due to the autapomorphy it possesses within the order of Psittaciformes. The genus demonstrates a unique physiology and biology, which is also observable in its biochemistry values (36). Consequently, high medians in HDL-C and LDL-C in this genus may not adequately compare to other psittacines. In the author’s avian practice *Eclectus* showed a prevalence and susceptibility for atherosclerosis similar to the genera *Psittacus* and *Amazona*. For the present study the total number of HDL-C and LDL-C measures in *Eclectus* parrots were only n = 43 compared to a much higher amount in amazons and African greys, so the results have to be interpreted with caution. More *Eclectus* focused studies will be needed to determine the scale of their susceptibility to atherosclerosis and the correlation between high HDL-C and LDL-C values. In the GLM with atherosclerosis as dependable variable, the only statistically significant genus related difference was recorded for African grey parrots in relation to amazon parrots. The genus *Psittacus* showed a higher predicted probability of developing atherosclerosis than the genus *Amazona*. For the other genera, no statement could be made, and while this may be due to the fact that the aforementioned genera had the largest sample sizes in this study, the result of the high atherosclerosis susceptibility of the genus *Psittacus* was in line with observations from previous studies (9, 37).

When studying both the LMM for HDL-C and LDL-C for their relation to atherosclerosis, a significant change in the predicted levels of LDL-C in presence of moderate to strong atherosclerosis was observed. However, in the GLM with atherosclerosis as a dependent variable, high levels of LDL-C or HDL-C were not significant predictors of atherosclerosis. This however may be due to the change of the character of the variable from categorial in the LLM into binary in the GLM. Diagnoses of light atherosclerosis were mixed together with moderate to strong atherosclerosis, and potentially observable effects for stronger atherosclerosis may have been obscured. Consequently, it can be assumed, that while there is a correlation between the presence of moderate to strong atherosclerosis and a high LDL-C value, a high LDL-C value itself cannot reliably predict or diagnose atherosclerosis. When looking at this result it makes sense, that a high LDL-C has to be interpreted as a potential risk factor and indicator for atherosclerosis, but not a definitive statement for the presence of atherosclerotic disease. Other than in a study conducted by Beaufrère, Cray (29), where a potential connection between atherosclerosis and a high HDL-C was presumed, our results indicate a correlation between atherosclerosis prevalence and increasing LDL-C, rather than HDL-C. HDL-C showed no statistically significant relation to atherosclerosis prevalence in the present study, contrary to our hypothesis. This result aligns with studies conducted in human medicine, where LDL-C has been identified as a major prognostic factor in the development and further evolution of atherosclerosis and cardiovascular disease. Naturally, the extrapolation of scientific findings from humans to birds has to be interpreted with caution due to the considerable species barrier, even though animal models play a crucial part in studying the pathomechanisms of atherosclerosis (18). In human medicine mendelian randomization studies and randomized controlled trials consistently demonstrated a log-linear relationship between the risk of atherosclerotic cardio-vascular diseases (ASCVD) and the absolute changes in plasma LDL-C (19, 20, 38–40). Biological and experimental evidence, in addition to the consistency among these studies, provides compelling evidence that LDL-C is causally associated with the risk of ASCVD in humans. Contrary to that, mendelian randomization studies about HDL-C, do not provide compelling evidence that HDL-C is causally associated with the risk of ASCVD (39, 41, 42). There is much rather an inverse association between HDL-C and the risk of ASCVD (20), which continues to be subject of discussion. In chickens, HDL deficiency syndrome did not result in increased susceptibility to atherosclerosis (43). In another study, chickens on an atherogenic diet had increased carotenoid, cholesterol and protein content in the LDL fractions, but not the HDL fractions (44). In reptiles, atherosclerosis is scarcely reported and consequently poorly understood (45). However, when present, LDL-C has been speculated to be the most prominent risk factor for its development (45, 46).

A study on lipoprotein cholesterol concentrations in 31 parrots of the genus *Amazona* (30) found no significant correlation between lipoprotein cholesterol concentrations and age (younger or older than 20 years), sex, and diet, apart from a difference in HDL-C concentration in male and female amazons, the latter which we were not able to confirm in our study. Our results further suggested that BCS in parrots seems to be connected to HDL levels, but not to LDL levels.

While we confirmed that there was no connection between HDL-C and diet, there appeared to be a significant correlation between LDL-C and diet, like it was hypothesized. Seed diets in comparison to pelleted or extruded diets have been shown to contain excess fat, low calcium to phosphorus ratios and many other deficiencies in vitamins and minerals (47–49). Birds fed a balanced diet of extrudates or pellets showed significantly lower LDL-C values than birds on a mixture of different foods or a pure seed diet. At the same time diet had an overall influence on the prevalence of atherosclerosis. Birds on a diet from categories 2 and 3, reliably displayed lower probability for the development of atherosclerosis as compared to the birds that were on a pure, unbalanced seed diet. Diet-induced atherosclerosis has been demonstrated experimentally in budgerigars (*Melopsittacus undulatus*) and Quaker parrots (*Myiopsitta monachus*) (28, 50). Quaker parrots fed a 1% cholesterol diet developed severe dyslipidemias with marked increases of TC and LDL-C and advanced atherosclerotic lesions in the aorta, brachiocephalic trunks, and coronary arteries within 4 months (28). TC by itself however has proven to be an inadequate marker for the diagnosis and development of atherosclerosis, and in clinical situations the single evaluation of TC in regards of diagnostics for atherosclerosis lacks significance. Birds have been observed with atherosclerosis, but normal TC values, however parrots with high TC and the absence of atherogenic changes have been commonly reported as well (51, 52). In addition, female parrots undergoing seasonal reproductive activity and vitellogenesis showed large scale elevations of TC and triglycerides among other biochemical parameters without vascular disease being present, further complicating the evaluation of TC (53, 54).

In this study, contrary to our hypothesis sex had no significant effect on neither HDL-C and LDL-C concentrations, nor on the prevalence of atherosclerosis. Other studies on lipoproteins in psittacines were inconsistent regarding sex differences for lipids too. While some also showed no difference (29), others marked elevated HDL-C levels in females compared to males (30) or higher LDL-C in females (55). Surprisingly, in this study, reproductive status at time of sample collection showed no effect on HDL-C and LDL-C values, apart from a significance (p-value of 0,01) for LDL-C for females that reportedly laid an egg within one week before or after sample collection (category 4). Due to the small number of observations in category 4, this significance is likely to be attributed to an instability of the model. If the significance was absent in a larger sample size, it could be due to the classification system that was used. Reproductive behaviour in itself is difficult to classify, and the time window of one week before and after egg laying in category 4 may have been too large and doesn’t adequately reflect the actual shifts in female lipid metabolism during vitellogenesis. However, in the event of a confirmation of this effect with a larger sample size, these changes would likely relate to the effects of oestrogens on the lipid metabolism, as well as protein and calcium metabolism in reproductively active females. Oestrogen promotes increased plasma TC, TG, calcium and protein levels (56). These changes may promote atherogenesis and may at least partially explain the association between reproductive tract disease and increased prevalence of advanced atherosclerotic lesions (9). For future studies another time window of three days should be used in a larger population, to reevaluate the effects of reproductive behaviour on HDL-C and LDL-C values. Breeding status was studied for its correlation with the presence of atherosclerosis as well, but showed no significant effect in that matter. While there was an expected correlation with age, from an evolutionary perspective it makes sense that seasonal reproductive behaviour would show no immediate effect on the presence of atherosclerosis, as this would be counterproductive to the continued existence of a species, if the majority of their individuals developed marked disease after reproducing multiple times.

The present study results showed, that age had no significant effect on HDL-C or LDL-C levels, which was in line with another study on its influence on the development of atherosclerosis (29). As atherosclerosis is characteristically a slowly progressing disease, it makes sense that with increasing age, there will be more individuals with the disease present. It is likely that, as in humans, all birds will eventually develop some degree of atherosclerotic disease with rising age, which may or may not be clinically distinct and actually cause symptoms. At that point the individual needs to be evaluated for other present risk factors and overall health status, so actual therapy and preventive measures can be applied.

There are some limitations within this study and the reference intervals established. A significant limitation is given due to the fact, that all parrots in the study were pets or breeding birds and originated from different places, where they were kept under different settings. While this variability renders the sample pool inhomogeneous, it concurrently offers a representative cross-section of bird species commonly encountered in daily avian practice. Though avoiding exercise and a sedentary lifestyle is another known risk factor for atherosclerosis in humans, the lack of a classifiable scale and unequal husbandry conditions made an evaluation impossible and for this reason was excluded from this study. The same is applicable for genetics. Genetic influences on serum lipoprotein levels in humans have been studied for many years now, and a reference study from as early as 1990 demonstrated that the proposed genetic locus accountable for LDL-C subfraction phenotypes eventually results in an atherogenic lipoprotein phenotype (57). Genetic disorders, like familial hypercholesterolemia, probably the most common monogenic dyslipidemia, can cause premature atherosclerosis and cardiovascular complications (58). However, while hypercholesterolemia is common in psittacine birds (54, 59), avian medicine lacks identified genetic disorders for parrots so far, and though some of the parrots in this study might be affected, they consequently fell under the radar.

Another limit is the method for determining the HDL-C and LDL-C values, which in this study were measured on a classic biochemistry analyser as direct measurement, rather than on a more accurate, but less available and high cost, high performance liquid chromatography (HPLC). HDL-C has been proven to be very accurate, when measured on a biochemistry analyser, and while LDL has been shown more difficult to properly measure (60), it at the moment is the most convenient way to obtain LDL-C values. It has been hypothesized that instead of LDL-C, non HDL-C should be used in clinical situations (60). The Friedewald formula is another commonly used equation and indirect method to calculate LDL-C values in humans, but has shown to be inaccurate for parrots, especially in lipemic samples (54, 60). In this study only HDL-C and LDL-C without their existing subclasses were evaluated. While today it is clear that specific subclasses of LDL-C, like the small-dense-LDL-C, are more atherogenic than others (61), pilot studies and base intervals for the main groups of HDL-C and LDL-C had to be performed first, before subfractions can be evaluated.

Because of lacking standard procedures, occurrence of atherosclerosis within the present study was diagnosed by various techniques. X-Ray has been shown to have a low sensitivity, when it comes to diagnosing atherosclerosis (6, 14), which is why only birds with severe detectable calcification of the great vessels were included. Endoscopy, though an invasive procedure, is a useful tool to identify various diseases in pet birds (62) and has been described to be used for the diagnosis of atherosclerosis in parrots before (7, 17, 33, 63). For this study, the large heart vessels and aorta descendens were evaluated. However, assessments have up until now never been cross checked with histologically based investigations. The results of this subjective evaluation may differ from the actual extend of the lesions. Further studies on the value of this invasive endoscopic method for diagnosing atherosclerosis will be needed in the future. The outcomes of post-mortem examinations were also used for diagnosis of atherosclerotic disease. Post-mortems provide the best results for identifying atherosclerotic lesions, and offer insight into the extent of the disease in retrospect (6, 9). Additionally. it allowed the subjective comparison of actual results with ante mortem diagnostics, which in this study came to the same conclusions.

In conclusion, the lipoprotein panel has the potential to become a screening tool for clinicians in the diagnosis of dyslipidemias. This study provides new insight on the value of HDL-C and LDL-C in parrots and provides reference intervals for many different genera of psittacines. Next to prominent genus related differences for HDL-C and LDL-C, the later has shown to be elevated in the presence of moderate to strong atherosclerotic disease and is influenced by diet. However, it is not a sole indicator of the presence of atherosclerotic disease and should rather be interpreted as a risk factor alongside variables like age and diet, which have also been shown to be associated with atherosclerosis. The clinician should therefore put special focus on lipoproteins, diet and genus, when it comes to the individual risk of a parrot developing atherosclerotic disease. However, further studies will be needed, to more indefinitely define the link between lipoproteins and atherosclerosis in parrots.

## Acknowledgments

We would like to acknowledge and thank the staff of Antech Augsburg, Germany (previously Synlab Augsburg) for their continued assistance and cooperation over the course of this study. No external funding or sponsoring was contributed to the development of this work.

## Author contribution

**Conceptualization:** Matthias Janeczek, Rüdiger Korbel, Friedrich Janeczek, Helen Alber, Helmut Küchenhoff, Monika Rinder.

**Data curation:** Matthias Janeczek, Friedrich Janeczek.

**Formal analysis:** Helen Alber, Helmut Küchenhoff.

**Investigation:** Matthias Janeczek, Monika Rinder.

**Resources:** Matthias Janeczek, Friedrich Janeczek, Rüdiger Korbel.

**Supervision:** Rüdiger Korbel.

**Writing – original draft:** Matthias Janeczek, Helen Alber, Monika Rinder.

**Writing – review & editing:** Matthias Janeczek, Rüdiger Korbel, Friedrich Janeczek, Helen Alber, Helmut Küchenhoff, Monika Rinder.

## Supporting Information

**S1 Table. β, confidence intervals (CI) and p-values for coefficient estimates of covariates in HDL-C model.** Reference categories are no atherosclerosis, *Amazona*, BCS of 3, male, fed according to category 1 and the reproductive status of category 1.

**S2 Table. β, confidence intervals (CI) and p-values for coefficient estimates of covariates in LDL-C model.** Reference categories are no atherosclerosis, *Amazona*, BCS of 3, male, fed according to category 1 and the reproductive status of category 1.

**S3 Table. β, confidence intervals (CI) and p-values for coefficient estimates of covariates in atherosclerosis model.** Reference categories are Amazona, BCS of 3, male, fed according to category 1 and the breeding status of category 1.

**S1 Fig. Flowchart of regression populations and their exclusion process of birds S2 Fig. Effect plot with confidence intervals for HDL-C and atherosclerosis.**

**S3 Fig. Effect plot with confidence intervals for LDL-C and BCS with outlier.**

**S4 Fig. Effect plot with confidence intervals for HDL-C and BCS without outlier.**

**S5 Fig. Mosaic plot of Age and atherosclerosis.** X-axis giving age in years in categories 1-10, 11-20, 21-25, 26-30 and >30; Y-axis indicating presence of atherosclerosis in yes and no.

**S6 Fig. Mosaic plot of diet and atherosclerosis.** X-axis giving diet in category 1 pure seed diet, category 2 mixed diet: seeds, table food, pellets/extrudates, category 3 pure pellets/extrudates; Y-axis indicating presence of atherosclerosis in yes and no.

**S1 File. R-Code and instructions for running.**

**S2 File. Dataset.**

**S3 File. Definition and categorization of investigated independent variables.**

